# Spawning Drivers of Female Atlantic Sturgeon (*Acipenser oxyrinchus oxyrinchus*) in the James River: A Fine-Scale Temporal Analysis

**DOI:** 10.64898/2026.05.21.726774

**Authors:** Matthew T. Balazik, Austin J. Draper, Greg C. Garman

**Affiliations:** Environmental Laboratory Coastal Ecology Team Army Engineer and Research Development Center 3909 Halls Ferry Rd. Vicksburg, MS 39180; School of Life Sciences and Sustainability, 1000 West Cary Street, Virginia Commonwealth University, Richmond, VA 23284

**Keywords:** Atlantic Sturgeon, sturgeon, spawning, resource management, endangered species

## Abstract

Fall-run Atlantic Sturgeon occupy riverine habitat for extensive periods of up to several months during periods of spawning. It is generally not feasible to eliminate potential anthropogenic stressors during the entire time Atlantic Sturgeon are on spawning habitat. If data were available to accurately predict when eggs and larvae are in the water column, potential impacts could be minimized for this relatively short timeframe, compared to the entire season of freshwater residency by adults. This research used acoustic telemetry data for adult female sturgeon, along with water temperature and discharge, to predict when females were likely releasing eggs versus merely staging in spawning habitat waiting to spawn. The descriptive and Bayesian model results predict that egg release is associated with water temperatures ranging from 20-26°C and pulses in river discharge, often confining egg release to a few days or weeks. By using weather data that predict relatively short periods of when egg release occurs versus longer periods that includes staging, resource managers can more feasibly collaborate with water usage groups to ameliorate egg/larvae survival.

## Introduction

Acipenseriformes, sturgeon and paddlefish, are a group of relict bony fishes with most species designated as threatened by the International Union for Conservation of Nature. There are eight species of sturgeon in the United States, all of which have state or federal protection. Major reasons for most historical sturgeon population declines included overfishing, degradation of spawning habitat, and spawning habitat exclusion due to dams [1,2]. Sturgeon have relatively long lifespans, delayed maturity, and intermittent spawning which extends the time needed for population recovery [1,3,4]. Understanding movement ecology, specifically of spawning behavior, for threatened and endangered sturgeon species is essential for effective conservation and restoration.

The anadromous Atlantic sturgeon (*Acipenser oxyrinchus oxyrinchus*) is federally listed as threatened or endangered within United States waters [5,6]. The reproductive success of sturgeon populations depends on precise timing and location of spawning events to optimize physicochemical conditions for early life stages. Significant efforts have been made to understand adult Atlantic Sturgeon migration and spawning site fidelity [7,8,9,10,11,12]. However, the fine-scale (i.e., days-weeks) timing of when spawning is occurring and eggs/larvae are in the water column and vulnerable to impingement/entrainment, versus adults occupying spawning habitat pre- and post-spawn, has not been thoroughly studied for Atlantic sturgeon.

The James River is the southernmost Atlantic sturgeon spawning tributary within the Chesapeake Bay and supports both spring and fall spawning Atlantic sturgeon populations [4,7,12,13,14]. Like other Atlantic slope rivers, the James River has been modified for a wide range of municipal and industrial uses, including water withdrawal, power generation, and shipping. Various portions of the upper tidal James River where fall spawning occurs have deep-draft shipping needs, water withdrawal for power generation and industrial cooling, and municipal drinking water which may negatively impact eggs and larvae. Similar types of hydromodification and environmental stressors have the potential to affect river-resident sturgeon across the species’ range [15].

Male and female fall Atlantic sturgeon enter the James River as early as April and typically stage in the lower (rkm 30-90) James for months before heading upstream in late summer/early fall [7]. Similar to the Hudson River [8], James River tagged males reach spawning habitats 2-5 weeks prior to tagged females and often stay at least two weeks after all tagged females have left. Some years individual males inhabit spawning areas 12 weeks of the year while residence times for females rarely surpasses four weeks [7,13].

If parameters of the critical phase of egg release and subsequent larval development can be predicted, adaptable planning beforehand may enable the reduction of potential negative anthropogenic impacts on egg/larvae survival. The goal of this study was to describe female spawning behavior to elucidate possible parameters that induce actual spawning and create a probabilistic model that predicts when Atlantic sturgeon eggs and subsequent larvae are likely to be present during the James River fall spawning season. By utilizing only female Atlantic sturgeon acoustic telemetry data and environmental variables, a model can support real-time management decisions aimed at reducing impacts on spawning success.

## Methods

All sampling and work conducted were approved by guidelines set by VCU’s Institutional Animal Care and Use Committee (#AD20127) and the National Marine Fisheries Service endangered species permit (#20314-01). VCU’s Institutional Animal Care and Use Committee permit was provided by VCU Animal Care and Use Program which is accredited by the International Association for Assessment and Accreditation of Laboratory Animal Care with continuous accreditation since August 3, 1966.

### Study Area

The James River, river mouth located at 36.98891, -76.30362, is the southern-most major tributary of the Chesapeake Bay (Fig 1). The James River is 696 km long and drains 26,164 km^2^. The tidal portion extends up to Richmond, VA located at rkm 155. River width varies between 0.7 and 7.1km up to rkm 120 and then narrows (range: 0.1 to 0.4 km). The federal navigation channel maintained by the U.S. Army Corps of Engineers runs from the river mouth at Hampton Roads, VA to rkm 150 and is maintained to a minimum depth of 7.6 m and a minimum width of 91.4 m.

**Fig 1.**
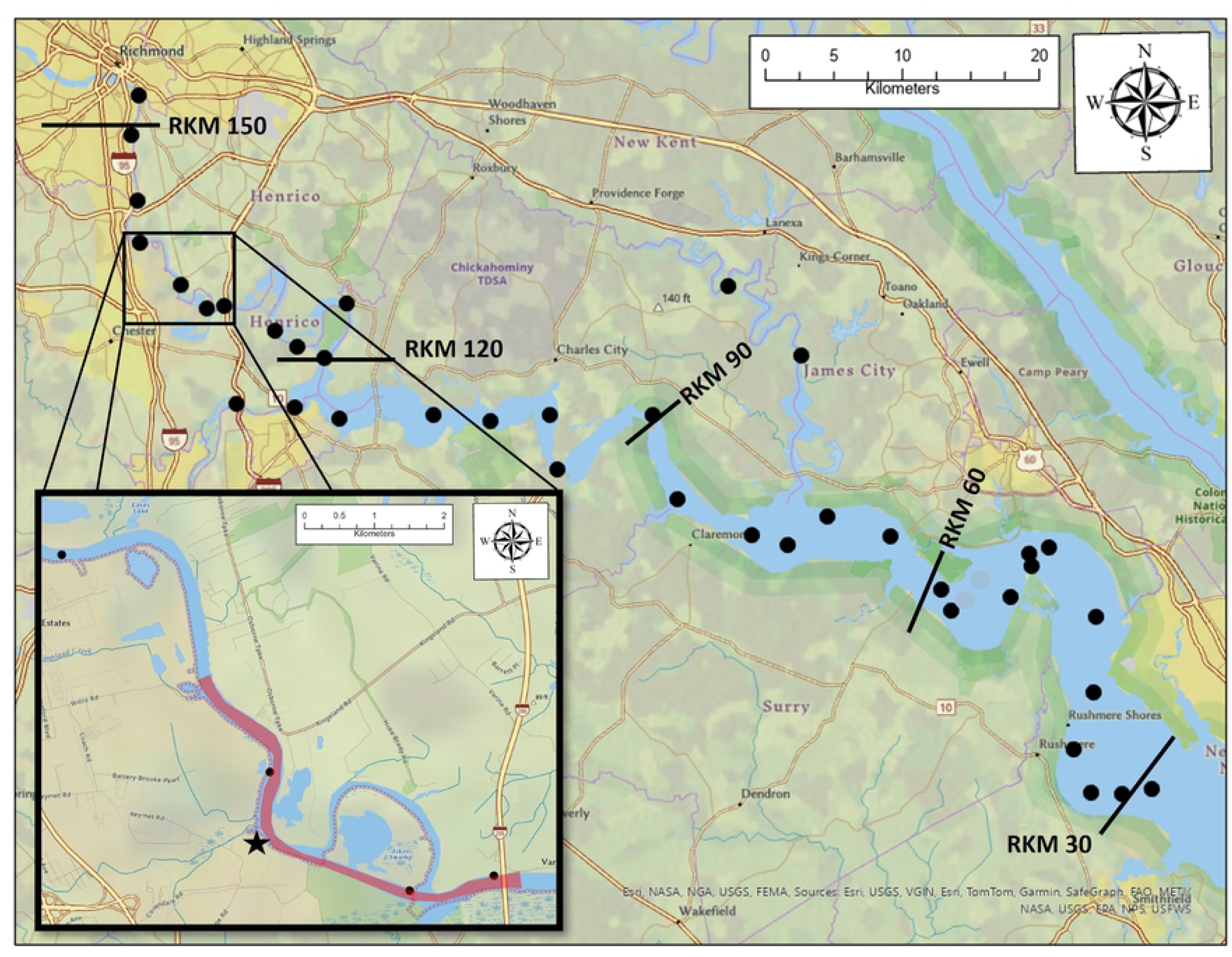
Map showing James River river kilometers and telemetry array (black dots). Note that some receivers were added, lost and not replaced during the 2020-2025 tracking period. The inset map shows the main fall Atlantic sturgeon spawning habitat from river kilometer 133-136 (highlighted red). The black star is the location of entrainment studies.

Between 2008 and 2025, gillnet sampling captured 940 unique adult fall-run Atlantic sturgeon, based on genetic analysis, in the James River [7,13,14]. Starting in 2014, sampling targeted pre-and post-spawn females staging around rkm 40, resulting in 28 telemetered females. Targeting pre-spawn fish staging at rkm 40 was discontinued in 2016 to reduce the potential impacts of capture on female spawning runs. While targeting males within 5km downstream of spawning habitat, eleven females were telemetered from 2017-2023 (Figs 1 & 2). All telemetry tags were surgically implanted using electronarcosis for anesthesia [16].

**Fig 2.**
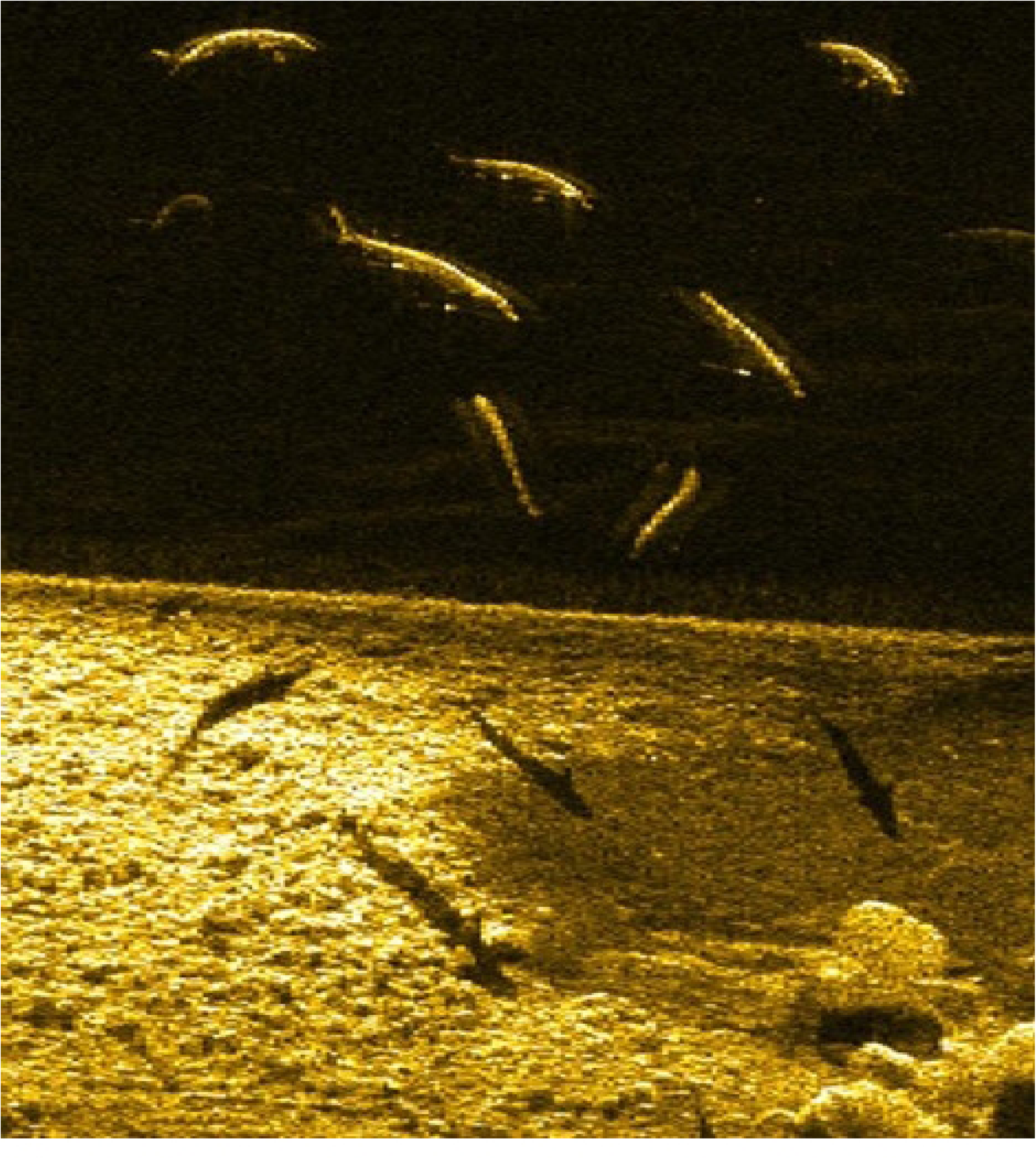
Example sidescan image of adult Atlantic sturgeon over cobble and rock habitat in the red highlighted area of Fig 1.

In 2017, the VCU James River telemetry receiver array was deployed to detect tags between rkm 30 and the fall line at rkm 155 (Fig 1). The array also provided coverage of major James River tidal tributaries including the Appomattox and Chickahominy Rivers (Fig 1). Based on our telemetry results, eggs/larvae collections [17] benthic habitat, and unpublished sidescan sonar data (Fig 2), the main spawning habitat in the James River is located at rkm 133-136. Occasionally, tagged females were detected at rkm 155 for a day or two but dropped back down and spent several days back at rkm 133-136 (Fig 1). During multiple years, Atlantic sturgeon have been observed spawning at the Appomattox River fall line but over 90% of females and males spend the entire spawning season in the mainstem James River.

James River post-spawn females are routinely captured at rkm 40 in early October/mid-November. A gravid female has never been captured during October/November downstream sampling efforts. While pre-spawn females are captured near spawning habitat each year, only two post-spawn females have been captured near spawning habitat. The low number of females upstream versus downstream is due to mesh size used to sample. Smaller gill net sizes, 25-33cm stretch mesh, are used near spawning habitat as the goal of the upstream sampling is to capture males and not females and females tend to be too big for 33cm mesh. When targeting females downstream, stretch mesh nets of 35-46 cm were used.

Low collections of post-spawn versus pre-spawn females near spawning habitat suggests females do not stay upstream after spawning. The best example of knowing the timing of a fall female spawning in the James River was in 2017 when a hydrated pre-spawn female was tagged near spawning grounds on October 3^rd^ and was later recaptured at rkm 40 on November 2^nd^. When recaptured, the ventral side was extremely concave which is a sign of a post-spawn female [18]. After being tagged near spawning grounds, the female remained at spawning habitat from October 3-5 when water temperatures were 22-23° C and then moved downstream to the post-spawn staging area at rkm 40. It is unknown how the impact of being captured and tagged modified the female’s spawning behavior. Based on the female’s ventral concave appearance and not returning to spawning habitat until 2020, three years is a common spawning interval for fall female James River Atlantic sturgeon [1]; it seems the female spawned during the October 3-5 timeframe and left immediately afterwards. Twelve pre-spawn and two post-spawn females were captured in the upper James between water temperatures 27-21°C and at 23°C, respectively (Fig 3).

**Figure 3.**
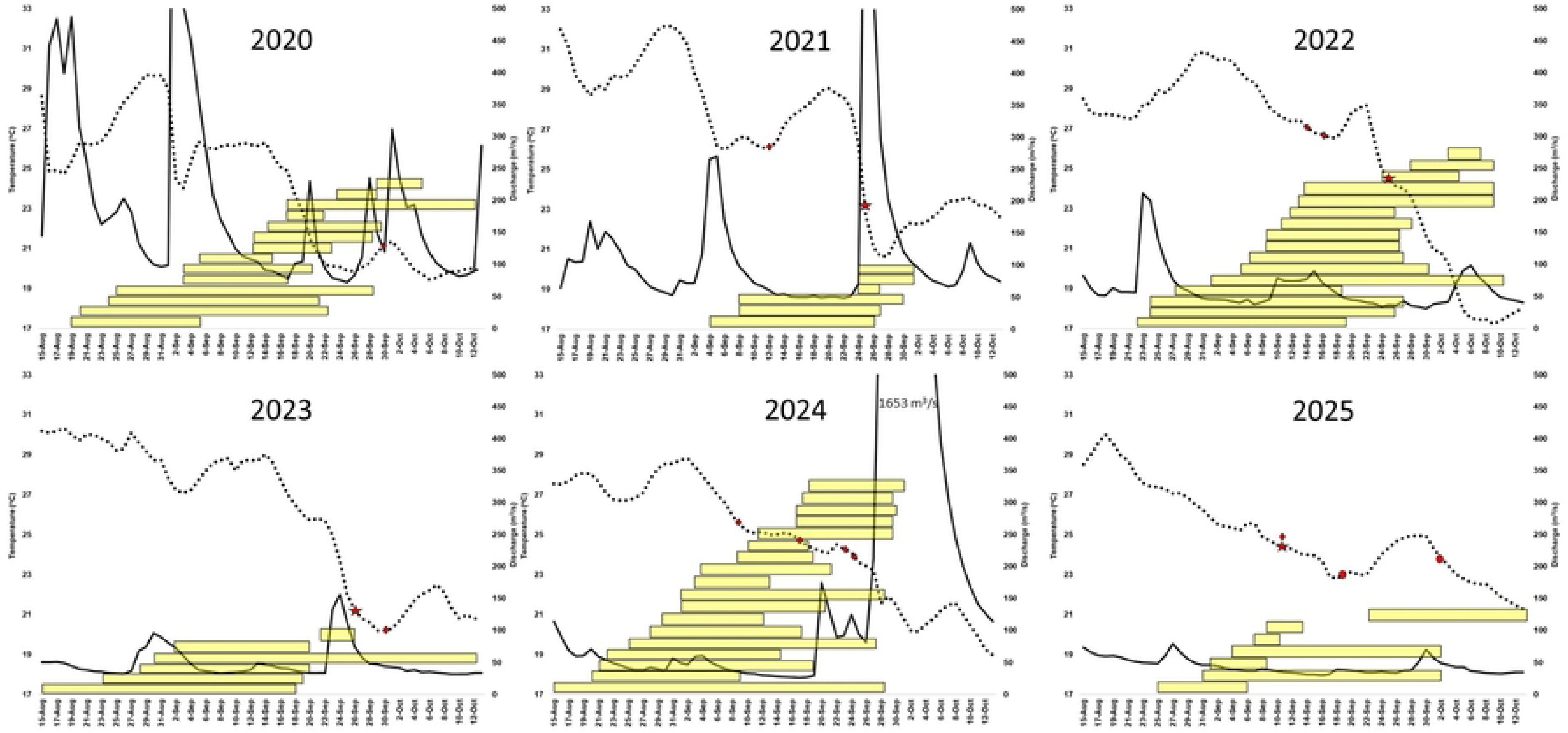
Multipanel plots showing when female Atlantic sturgeon first reached and last left spawning habitat each year (yellow horizontal bars). The black dashed line is water temperature at spawning habitat and the black solid line is river discharge at Richmond. The red star in 2021, 2022, 2023, and 2025 is when eggs were collected at rkm 134. The red diamonds and red circles are female pre- and post-spawn collections, respectively.

Bruch and Binkoski [19] using direct observations in Wisconsin’s Lake Winnebago system, described female lake sturgeon (*A. fulvescens*) typically take 8-12 hr but up to 18 hr from the onset of egg expulsion to leaving spawning habitat for the year. Because oocyte development typically takes at least a year, female Atlantic sturgeon are limited to one spawning event a season and likely leave immediately after spawning like lake sturgeon. Unlike females, male Atlantic sturgeon generate sperm throughout the spawning season, enabling them to spawn multiple times during the season [13]. Female egg release happens once a season and likely takes less than a day upon onset with the female leaving immediately afterwards; therefore, using when a female leaves spawning habitat for the year is a logical indicator of when an unaltered, *i.e*., not captured during the spawning season, female actually spawned versus roaming around spawning grounds waiting to spawn.

The goal of this study was to provide fine-scale timing of when female Atlantic sturgeon exhibit spawning behaviors associated with egg release (herein referred to as assumed spawn) in the James River along with water parameters that may induce and predict assumed spawning. One strategy plotted water parameters and female telemetry data from returning tagged females that had not been captured during the current migration season. A second method was a classification scheme based on daily acoustic telemetry data from female Atlantic sturgeon. Each day was assigned a behavior classification using telemetry data per transmitter per day. Data processing in R grouped detections by transmitter and datetime, using only the final detection of the day to summarize behavior. A female’s status was determined to be pre-spawn, assumed spawning, or post-spawn; however, post-spawn was not factored into the model. These classifications allowed binary logistic regression models (assumed spawn vs. pre-spawn) using environmental predictors to predict assumed spawning (Table 1).

**Table 1.** Classification scheme mapping daily behavioral observations to binary model outputs used in logistic regression. Spawning-related behaviors were assigned a value of 1 (“assumed spawn”), and non-spawning behaviors were assigned a value of 0 (“pre-spawn”).

| Classified action for each day | Numerical output of logistic regression model (0 = pre-spawn, 1 = assumed spawn) |
| --- | --- |
| Pre-spawn | 0 |
| Assumed spawn | 1 |

To be classified as an assumed spawning female, the fish must have made at least one movement upstream to spawning habitat. Based on sidescan sonar surveys, water quality, and telemetry data, spawning for fall Atlantic sturgeon is above rkm 118 where the Appomattox and James Rivers split [7] (Fig 2). This rule was implemented as some fall females stage in the lower James River during the summer and fall but do not move upstream of rkm 70. These staging females leave the James River in the fall and typically return the subsequent year to fall spawning habitat.

A female’s assumed spawning date was designated to be the day when the female last left spawning habitat for the year. A pre-spawn designation are days prior to the assumed spawning day. For example, in 2022 tag 16703 moved up to spawning habitat in late August as water temperatures were decreasing. Water temperatures increased to 31°C and tag 16703 dropped downstream to rkm 90. Water temperatures decreased from 30°C to 27°C during September 7^th^ to 10^th^ and tag 16703 moved back upstream to spawning habitat on September 11^th^, staying until the evening of September 16^th^. The fish moved downstream on the 16^th^, not returning to spawning habitat the remainder of the year. Days prior to September 16^th^ would be classified as pre-spawn, September 16^th^ would be considered spawning, and days after September 16^th^ would be considered post-spawn (Table 1).

Temperature is seen as the major driver for sturgeon spawning [1,19,11, 20]. River discharge being mostly associated with rain events is a major driver for water temperature fluctuations throughout the upper-tidal James and Appomattox Rivers. Water temperature and discharge were utilized by both strategies to elucidate fine-scale female assumed spawning behavior. All water temperatures used for this project were taken roughly 30cm from the bottom in the middle of the main spawning habitat of the mainstem James River, +37.39395, -77.38383. Water temperature was taken at 15min intervals and averaged for the entire day. Average daily James River discharge values were provided by USGS Gage 02037500 which is roughly 28 rkm upstream of the main spawning area.

Plotting water temperature, discharge, and female telemetry data was one method used to estimate female spawning trends. A Bayesian mixed-effects binomial logistic regression model was implemented using the rstanarm package [21] in RStudio to estimate assumed spawning timing. The model estimated the probability of an assumed spawning event as a function of temperature and discharge, with (in some model specifications) a random intercept for transmitter ID to account for repeated observations and individual-level variability.

Several alternative models (Table 2) were compared based on predictive performance using classification metrics derived from posterior predictions. For each model, the confusion matrix, overall accuracy, sensitivity (true positive rate), and specificity (true negative rate). The model with the best balance of sensitivity and overall accuracy was selected for interpretation and inference. All models were fit using the rstanarm package [21] in R.

**Table 2.** Summary of candidate models evaluated for predicting spawning probability. The asterisk (*) denotes the final model selected for analysis based on predictive performance metrics.

| Model | Description |
| --- | --- |
| Temp only | Includes only temperature as a fixed effect predictor of spawn probability. |
| Temp + Discharge | Includes both temperature and flow as fixed effect predictors. |
| *Temp + Discharge + (1 \| Transmitter) | Adds a random intercept for Transmitter ID to account for repeated measurements and individual-level variability. |
| Temp * Discharge + (1 \| Transmitter) | Added an interaction between temperature and discharge |
| 3-day lag Temp + Discharge + (1 \| Transmitter) | Replaces current temperature with a 3-day lagged temperature variable to capture delayed effects of environmental conditions, while including flow and a random intercept for Transmitter ID. |
| 3-day lag Temp * Discharge + (1 \| Transmitter) | Added an interaction between 3-day lag temperature and discharge |

Weakly informative priors were used because the strength of the relationship between temperature and discharge on the assumed spawning movements of Atlantic sturgeon is unknown and the model needed to be mostly informed by data given a lack of prior knowledge. For fixed effects (temperature and discharge) and random intercept term, (Transmitter) normally distributed prior distributions with a mean of 0 and a standard deviation of 2.5 was used.

The model was fit using 4 Markov Chain Monte Carlo (MCMC) simulations to assess convergence across independent sampling runs. Each chain was run for 10,000 iterations, including 5,000 warm-up (burn-in) iterations and 5,000 post-warm-up iterations used for inference. Model checking and diagnostics were done using the functions in Table 2 from the bayesplot package [22] in RStudio. Lastly, the Bayesrules package [23] in RStudio was used to assess the model’s posterior classification quality, including creating confusion matrices and calculating accuracy rates.

| Function | What It Checks |
| --- | --- |
| **mcmc_trace()** | Convergence & mixing of chains |
| **mcmc_dens_overlay()** | Same posterior across chains |
| **mcmc_acf()** | Autocorrelation in sampling |
| **neff_ratio()** | Sampling efficiency |
| **rhat()** | Convergence across chains |

Finally, simulations of how assumed spawning behavior would likely respond under a variety of water temperatures and discharges were performed by generating predicted probabilities from the model. Using 20,000 draws from the posterior distribution, the probability of assumed spawning under various conditions were simulated and visualized for comparison. These simulations were designed to explore how variations in temperature and discharge could instantaneously influence assumed spawning movements.

## Results

From 2020-2025, twenty-seven different females made 67 migrations to spawning habitat in the James and Appomattox Rivers and later left the system (Table 3, Fig 3). Fifty-five migrations inhabited only James River spawning habitat while five remained entirely within the Appomattox River. The remaining 12 migrations spent time in both systems but all ended up leaving the upstream areas after moving to James River spawning habitat, suggesting the fish spawned in the James River. There was one instance when a female returned to spawning habitats in two consecutive years. There were two instances when a female tag was lost in the upper James River during the spawning season and never relocated. Both tags still had years of battery life. Since the tags were never detected leaving spawning habitat for those two runs, the two runs were not used in analyses.

**Table 3.** Overview of various spawning parameters based on telemetry and egg collection data. All water temperatures are in Celsius. Arrival date is when the first telemetry tagged female arrived at spawning habitat each year along with the water temperature of that day. The first and last spawn dates are when the earliest and latest a telemetered female left spawning habitat each year. Days at spawning habitat lists the minimum, median, and maximum number of days from when a female first reached and left spawning habitat each year. Spawning water temperature range is the water temperatures spawning occurred based on telemetry data.

| Year | Number of females | Arrival date (water temp) | First spawn/last spawn (days) | Days at spawning habitat min/median/max | Spawning water temp range | Egg collections (water temp) |
| --- | --- | --- | --- | --- | --- | --- |
| 2020 | 14 | 19-Aug (25) | 16-Sep/12-Oct (27) | 3/15.5/34 | 26-20 | NA |
| 2021 | 6 | 4-Sept (26) | 25-Sep/1-Oct (7) | 4/13/22 | 23-21 | 25-Sept (23) |
| 2022 | 16 | 22-Aug (27) | 18-Sep/10-Oct (23) | 4/21/40 | 27-17 | 26-Sept (25) |
| 2023 | 8 | 15-Aug (30) | 23-Sep/12-Oct (18) | 6/29/43 | 26-21 | 26-Sept (21) |
| 2024 | 18 | 16-Aug (27) | 9-Sep/30-Sep (22) | 7/17.5/41 | 26-21 | NA |
| 2025 | 7 | 25-Aug (27) | 6-Sep/13 Oct (38) | 4/12/32 | 26-21 | 10-Sept (25) |

Individual females spent 3-43 days at spawning areas before leaving for the year. The number of days elapsed between the first and last female leaving spawning habitat ranged from seven in 2021 to 38 in 2025 (Table 3, Fig 3). Based on plotted telemetry and water quality parameters, decreasing water temperature seems to be a major driver influencing the timing of female assumed spawning in the James River, with discharge having some influence on water temperature. Females typically start arriving at spawning habitat in mid/late August when water temperature falls below 29°C (Table 3, Fig 3). Water temperatures in 2021 were relatively warm and tagged females did not arrive until early September when water temperature fell below 29°C. Females left spawning areas from 17-27°C with most (93%) spawning between 20-26°C (Fig 4). The only year females left spawning habitat below 20°C was 2022 when water temperature fell steadily due to cooler air temperatures and no rain events (Fig 3). Females that stayed relatively long arrived right when water temperatures fell below 29°C in mid-August but remained above 26°C for weeks as in 2022 and 2023.

**Figure 4.**
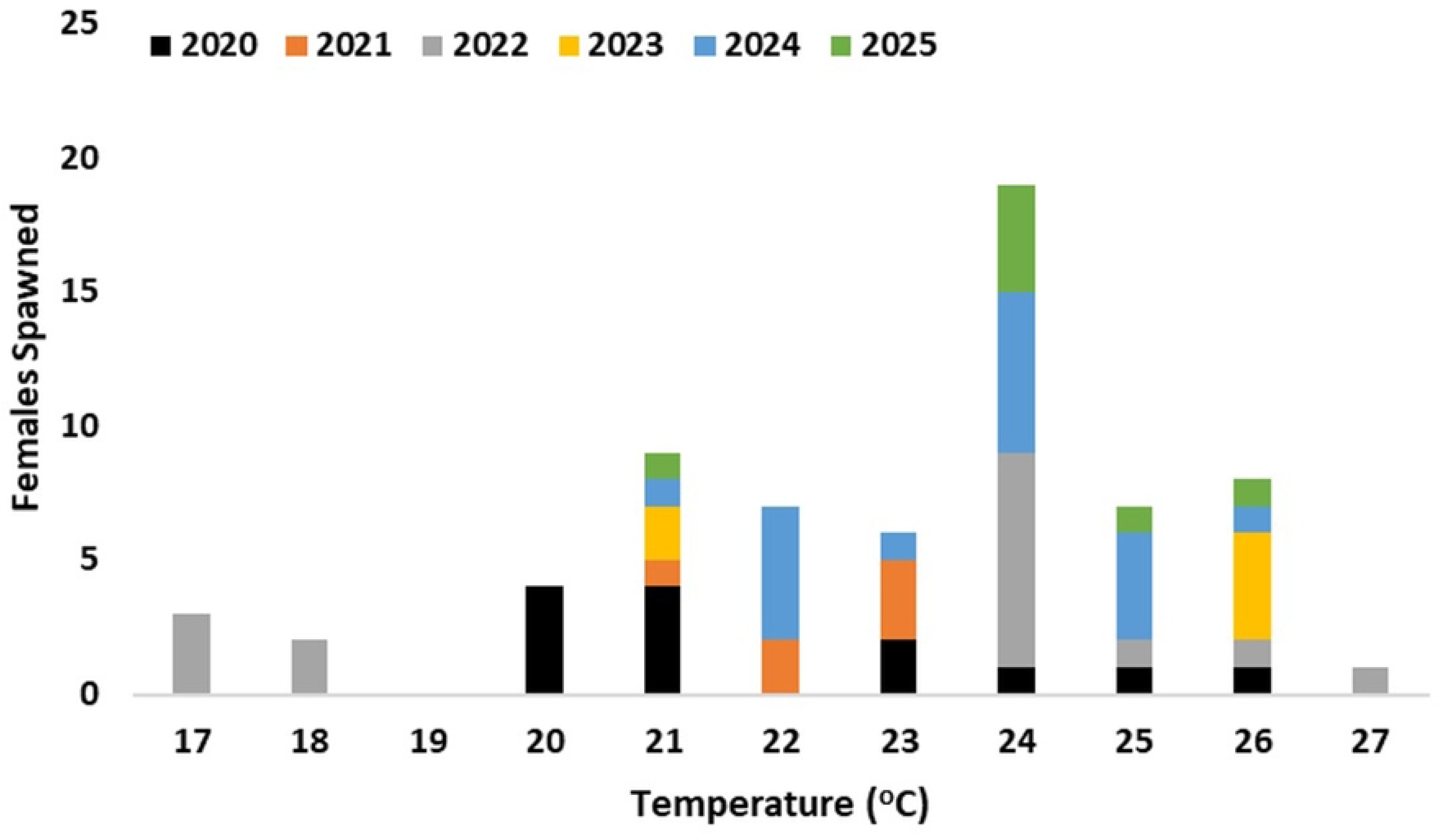
Water temperatures color coded by year for when females left spawning habitat for the year.

Once water temperatures decreased to 26°C, any increase in discharge with a coinciding fall in water temperature (2021, 2023, 2024) seemed to induce staging females to exhibit assumed spawning behavior. In 2022, there was no notable increase in discharge driving a decrease in water temperature. Rather an atypical fall in air temperature reduced water temperature from 28°C to 21°C over a nine-day period when 50% of females exhibited assumed spawning behavior within three days of each other. In 2025 there was a gradual decline from 26°C to 21°C and no notable discharge events which may be why that year had the widest assumed spawning season even though relatively few tagged females returned (Fig 3).

### Model diagnostics

Visual inspection of the trace plots, density overlay plots, and autocorrelation plots indicated good convergence and adequate mixing, suggesting that the MCMC sampler successfully explored the posterior distributions.

### Model evaluation

A low cutoff value of 0.05 was used to capture more assumed spawning events (increase sensitivity), which maintained strong overall accuracy, while keeping Type I error acceptably low given the model’s high specificity. The best-performing model was **Temp * Discharge + (1 | Transmitter)**, with an overall accuracy of 91% and sensitivity of 54.5% (Table 4). This means the classification cutoff value of 0.05 correctly predicted 54.5% of actual assumed spawning events and 91.9% of pre-spawning events. The primary model included a random intercept for transmitter ID, which improved performance relative to simpler models that excluded this term. The simpler models showed reduced predictive accuracy, underscoring the value of accounting for individual variation.

**Table 4.** Models fitted and resulting summaries of the model’s posterior classification quality. The model discussed in this paper is bolded. Model outputs were classified as spawn vs pre-spawn if the model prediction was over 5% likelihood of a spawn.

| Model variables | Sensitivity | Specificity | Overall accuracy |
| --- | --- | --- | --- |
| Temp + (1 \| Transmitter) | 0.415 | 0.933 | 0.897 |
| Temp + Discharge + (1 \| Transmitter) | 0.484 | 0.933 | 0.901 |
| **Temp \* Discharge + (1 \| Transmitter)** | **0.545** | **0.919** | **0.91** |
| 3-day lag Temp + Discharge + (1 \| Transmitter) | 0.915 | 0.711 | 0.725 |
| 3-day lag Temp * Discharge + (1 \| Transmitter) | 0.915 | 0.715 | 0.729 |

The confusion matrix and posterior classification summaries show that the classifications were correctly predicted whether a fish is in the spawning grounds 91% of the time (Tables 5-6). Specifically, the model demonstrates high specificity (92%) in identifying non-spawning events (Table 4). However, model sensitivity is lower, with only 54% of predicted assumed spawning movements corresponding to actual spawning events.

**Table 5.** Confusion matrix comparing model predictions to observed outcomes. Predictions were classified using a 0.5 probability threshold: predicted probabilities > 0.05 were labeled as “Spawn,” and those ≤ 0.05 were labeled as “Pre-spawn.” Each cell (a–d) is color coded to show whether model predictions correctly identified observed outcomes.

| Action | Pre-spawn<br>(from data) | Assumed spawn<br>(from data) |
| --- | --- | --- |
| No spawn (model prediction less than 50% probability) | 2406<br>(a) | 211<br>(b) |
| Spawn (model prediction over 50% probability) | 32<br>(c) | 34<br>(d) |

**Table 6.** Posterior estimates from the final Bayesian logistic regression model predicting spawning probability. Estimates are presented on the log-odds scale and include the mean, 80% credible intervals, and standard errors for each fixed effect term.

| Term | Estimate | 80% credible interval | Standard error |
| --- | --- | --- | --- |
| Temp | -0.449 | [-0.543, -0.365] | 0.0696 |
| Discharge | 0.0118 | [0.00260, 0.0213] | 0.00723 |
| Temp:Discharge | -0.000433 | [-0.000839, -0.0000451] | 0.000307 |

### Parameter evaluation

Parameter estimates from the final model are presented in Table 6. The temperature parameter had a mean estimate of –0.449, with an 80% credible interval of [–0.543, –0.365], entirely below zero. This indicates a strong negative relationship between temperature and assumed spawning probability: as temperature decreases, the likelihood of assumed spawning increases. The effect size of this relationship on the log-odds scale can be calculated using the equation shown in Equation 1. Specifically, for each 1°C decrease in temperature, the odds of assumed spawning increases by a factor of approximately 1.57, calculated as 1/exp(–0.449) = 1.57.

Equation 1. Calculation of the inverse odds ratio.

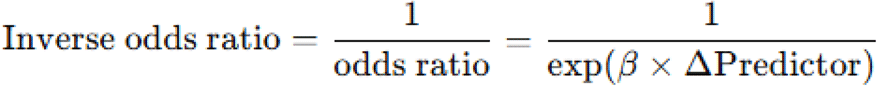

The discharge parameter had an 80% credible interval of [0.0026, 0.0213] (Table 6), suggesting that its main effect was not statistically significant on its own. In contrast, the interaction between temperature and discharge was significant, with a mean estimate of –0.0004 and an 80% credible interval of [–0.0008, –0.000005]. This interaction is visualized in Fig 5, which shows that at higher flow rates, a 5°C decrease in temperature (from 25°C to 20°C) leads to a substantially greater increase in the probability of spawning. For instance, at 20°C, the predicted probability of assumed spawning is nearly 90% under maximum observed flow conditions (1317 m³/s), compared to approximately 20% under minimum flow conditions (27 m³/s). These results indicate that the influence of temperature on assumed spawning probability is strongly moderated by discharge.

**Figure 5.**
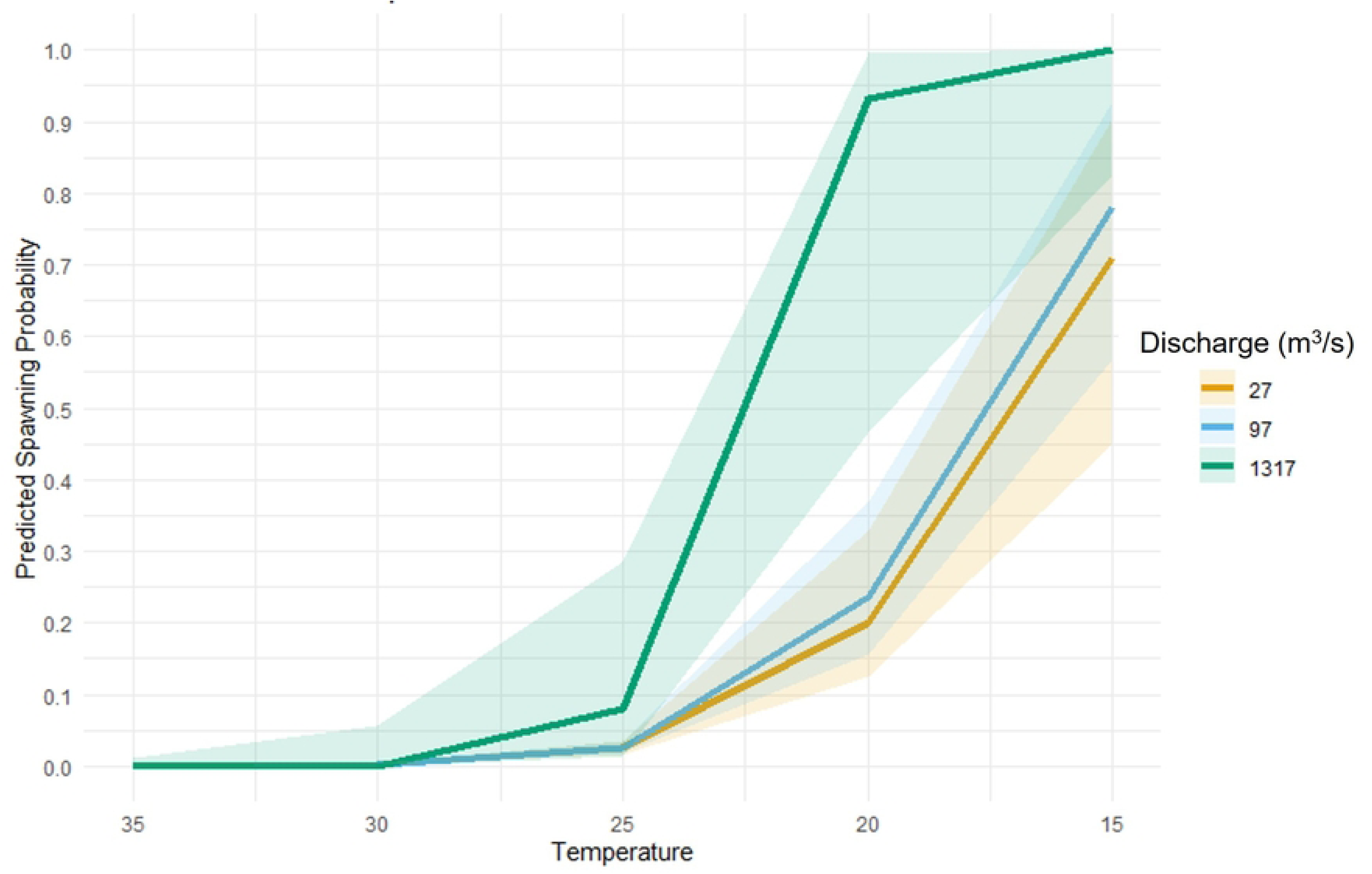
Predicted spawn probabilities at varying temperatures and flows. The y-axis represents the modeled predicted spawn probability and the x-axis is temperature. Flow is grouped by the minimum (orange), mean (blue), and max (green) of all m3/s values found in the 2020-2026 James River dataset.

## Discussion

These results are based on the assumption that all females spawned before leaving habitat and did not abort spawning due to some unknown factor anytime during the spawning season. It is possible the five female outliers that stayed late in 2022 did not spawn when water temperatures were 17-18°C but left and held their eggs for the following year or reabsorbed them. Telemetry data show that males were around the females when water temperature was 17°C, the females did not return the following year, and four returned to spawning habitat in 2024 while the fifth has not returned to James River. Furthermore, a gravid female has never been captured at the downstream staging area in October/November. Based on the overall results from the outliers, it is hypothesized they did spawn and fall spawning may occur in water temperatures at 17°C.

Telemetry and water quality data show that Atlantic sturgeon fall spawning likely occurs between 20-26°C water temperatures with most assumed spawning at 24°C (Figs 3 & 4). This brackets the 24.3-25.3°C temperature range that eggs were captured from natural spawning in the Roanoke River [10]. Discharge events that drive temperatures down relatively quickly through the spawning temperature range seem to cause most females to exhibit spawn behavior in a relatively short time period. The assumed spawning window seems to increase if there is no spike in discharge and water temperatures gradually decrease.

A better understanding regarding spawning patterns due to environmental conditions may elucidate trends in juvenile survival. By incorporating age 0-2y catch-per-unit-effort information, catch trends may show whether relatively short spawning seasons when temperatures fall relatively quicker from 26 to 20°C have better success then gradual decreases or vice versa. If it is shown that short spawning periods have reduced spawning success, climate trends of warmer summers and strong storm events causing rapid drops in temperatures may be problematic for spawning success. If a situation is found that shows increased spawning success, the pattern could possibly be mimicked by already installed water controls structures such as dams.

The overall classification accuracy of 91% and high specificity (92%) indicate that the model reliably distinguishes between assumed spawning and non-spawning events. However, the relatively low sensitivity (55%) suggests that many true assumed spawning events are not being captured. This limitation is expected, given that spawning is a relatively rare behavior and fish may spend extended time in the lower river before making brief excursions to spawning grounds (Fig 3). As such, the data are weighted more heavily toward non-spawning observations, influencing the model’s ability to detect spawning. Despite this, a 55% probability of correctly identifying an assumed spawning event remains valuable from a management perspective. When the model indicates spawning activity, it can serve as an early warning or decision-making tool, providing timely evidence for conservation or monitoring actions.

This approach also has implications for reducing anthropogenic threats that coincide with assumed spawning. Vessel strikes, water withdrawals, and egg/larvae entrainment into industrial facilities are possible threats to early life stages of Atlantic sturgeon [3]. It is impractical to stop all anthropogenic disturbances for when adult Atlantic sturgeon are near spawning habitat which can be up to 12 weeks. The ability to forecast more precise spawning behavior windows down to a few days enables targeted, adaptive mitigation strategies. For example, if conditions are predicted to reach optimal spawning thresholds (e.g., temperature <24°C and stable or declining discharge after a spike), managers could temporarily reduce water withdrawals or modify construction, shipping, or dredge schedules in critical river segments to lessen potential negative impacts to spawning behavior and reduce egg and larval loss.

This study demonstrates that fall run Atlantic sturgeon spawning behavior in the James River can be modeled and probabilistically predicted using environmental variables, particularly water temperature and discharge. By integrating acoustic telemetry with biological knowledge of Atlantic sturgeon behavior and spawning ecology, a classification scheme was successfully developed that enables the prediction of when females likely spawn and when their eggs and larvae are likely to be present in the system. Traditionally, direct observations from capture data or egg sampling have been used to infer spawning timing [1,3,10,18], but with technological advancements and integration of expert knowledge, periods of egg release may be predicted by looking at weather forecasts. The information from this study allows researchers and managers to estimate spawning activity at a fine temporal scale using water parameters. The use of expert-informed behavioral classifications and environmental cues fills a significant knowledge gap and represents a promising tool for endangered species management. While there are subtle differences, the water quality drivers that induce spawning behavior in the James River are likely germane to other fall spawning populations.

## Acknowledgements

We would like to thank Greg Garman (Virginia Commonwealth University), Peter Sturke and Stephen Dwyer (Dominion Energy), Douglas Clarke (Clarke Environmental LLC) and Martin Balazik, Thiwaporn Balazik, George Trice, and Charles Frederickson (field volunteers) for assisting with this project. This manuscript represents VCU Rice Rivers Center publication 104. The authors declare that they have no conflict of interest.

## Notes

### Competing Interest Statement

The authors have declared no competing interest.

## References

1. Hilton E, Kynard B, Balazik M, Horodysky A, Dillman C. Review of the biology, fisheries, and conservation status of the Atlantic Sturgeon, (*Acipenser oxyrinchus oxyrinchus* Mitchill, 1815). J Appl Ichthyo. 2016; 32: 30–66. 10.1111/jai.13242

2. Lotze H, Lenihan H, Bourque B, Bradbury R, Cooke R, et al. Depletion, degradation, and recovery potential of estuaries and coastal seas. Science. 2006 Jun 23; 312 (5781): 1806–1809. 10.1126/science.1128035

3. Atlantic Sturgeon Status Review Team. Status review of Atlantic sturgeon *(Acipenser oxyrinchus oxyrinchus)*. Report to National Marine Fisheries Service, Northeast Regional Office. 2007 Feb 23. 174 p. Available from: https://repository.library.noaa.gov/view/noaa/16197

4. Przelomska NAS, Balazik MT, Lin AT, Reeder-Myers LA, Rick TC, Kistler L. Archaeogenomic analysis of Chesapeake Atlantic sturgeon illustrates shaping of its populations in recovery from severe overexploitation. Proc Biol Sci. 2024 Oct 9; 291(2032): 20241145. 10.1098/rspb.2024.1145

5. Office of the Federal Register. Endangered and threatened wildlife and plants; threatened and endangered status for distinct population segments of Atlantic sturgeon in the northeast region. Fed Regist 2012a. 2012 Feb 6; 77 (24): 5880–5912. 50 C.F.R. Parts 223 and 224.

6. Office of the Federal Register. Endangered and threatened wildlife and plants; threatened and endangered status for distinct population segments of Atlantic sturgeon in the northeast region. Fed Regist 2012b. 2012 Feb 6; 77 (24): 5880–5912.

7. Balazik MT, Musick JA. Dual annual spawning races in Atlantic sturgeon. PLoS One. 2018 May 28; 10(5): e0128234. 10.1371/journal.pone.0128234

8. Breece M, Higgs A, Fox D. Spawning intervals, timing, and riverine habitat use of adult Atlantic sturgeon in the Hudson River. Trans Am Fish Soc. 2021 Feb 22; 150(4): 528–537. 10.1002/tafs.10304

9. Farrae D, Post W, Darden T. Genetic characterization of Atlantic sturgeon, *Acipenser oxyrinchus oxyrinchus*, in the Edisto River, South Carolina and identification of genetically discrete fall and spring spawning. Conserv Genet. 2017 Jan 24; 18: 813–823. 10.1007/s10592-017-0929-7

10. Smith J, Flowers H, Hightower J. Fall spawning of Atlantic sturgeon in the Roanoke River, North Carolina. Trans Am Fish Soc. 2015 Jan; 144(1): 48–54. 10.1080/00028487.2014.965344

11. Vine J, Holbrook S, Post W, Peoples B. Identifying environmental cues for Atlantic sturgeon and shortnose sturgeon spawning migrations in the Savannah River. Trans Am Fish Soc. 2019 May; 148(3): 671–681. 10.1002/tafs.10163

12. White S, Kazyak D, Darden T, Farrae D, Lubinski B, Johnson R, et al. Establishment of a microsatellite genetic baseline for North American Atlantic sturgeon (*Acipenser o. oxyrinchus*) and range-wide analysis of population genetics. Conserv Genet. 2021 Aug 02; 22: 977–992. 10.1007/s10592-021-01390-x

13. Balazik MT, Garman GC, Van Eenennaam JP, Mohler J, Woods III LC. Empirical evidence of fall spawning by Atlantic sturgeon in the James River, Virginia Trans Am Fish Soc. 2012 Nov; 141(6): 1465–1471. 10.1080/00028487.2012.703157

14. Balazik MT, Farrae DJ, Darden TL, Garman GC. Genetic differentiation of spring-spawning and fall-spawning male Atlantic sturgeon in the James River, Virginia. PLoS One. 2017 Jul 7; 12(7): e0179661. 10.1371/journal.pone.0179661

15. Reine K, Clarke D, Balazik M, O’Haire S, Dickerson C, Frederickson C, et al. Assessing impacts of navigation dredging on Atlantic sturgeon (Acipenser oxyrinchus). Vicksburg (MS): U.S. Army Engineer Research and Development Center; 2014 Nov. Technical Report No.: ERDC/EL TR-14–12. 10.13140/RG.2.1.5064.6565

16. Balazik MT. Capture and brief invasive procedures using electronarcosis does not appear to affect postrelease habits in male Atlantic sturgeon during the spawning season. N Am J Fish Manag. 2015 Apr 15; 35(2): 398–402. 10.1080/02755947.2015.1011358

17. Incidental Take Permit to Virginia Electric and Power Company, D.B.A. Dominion Virginia Power. NOAA Notice NMFS. 2025. Available from: https://www.fisheries.noaa.gov/action/incidental-take-permit-virginia-electric-and-power-company-dba-dominion-virginia-power

18. Ryder JA. The sturgeons and sturgeon industries of the eastern coast of the United States, with an account of experimental bearing upon sturgeon culture. Bulletin of the United States Fish Commission Vol. VIII, for 1888, 231–328.

19. Bruch RM, Binkowski FP. Spawning behavior of lake sturgeon (*Acipenser fulvescens*). J Appl Ichthyol. 2002 Dec 17; 18: 570–579. 10.1046/j.1439-0426.2002.00421.x

20. Paragamian V, Kruse G, Wakkinen V. Spawning habitat of Kootenai River white sturgeon, post Libby Dam. N Am J Fish Manag. 2001 Feb; 21(1): 22–33. 10.1577/1548-8675(2001)021<0022:SHOKRW>2.0.CO;2

21. Goodrich B, Gabry J, Ali I, Brilleman S. rstanarm: Bayesian applied regression modeling via Stan. R package version 2.32.1. 2024. Available from: https://mc-stan.org/rstanarm/

22. Gabry J, Mahr T. bayesplot: Plotting for Bayesian models. R package version 1.13.0.9000. 2025. Available from: https://mc-stan.org/bayesplot/

23. Dogucu M, Johnson A, Ott M. bayesrules: Datasets and supplemental functions from Bayes Rules! Book. R package version 0.0.2.9000. Available from: https://github.com/bayes-rules/bayesrules

